# PCP-dependent polarity propagation across neuronal columns in the *Drosophila* medulla

**DOI:** 10.64898/2026.05.26.727799

**Authors:** Takumi Morita, Haruka Yasuda, Jiro Osaka, Makoto Sato, Takashi Suzuki

## Abstract

Planar cell polarity (PCP) signaling establishes coordinated tissue polarity through asymmetric localization of core PCP factors and propagation of polarity information between neighboring cells. While PCP mechanisms are well characterized in continuous epithelial tissues, whether PCP-dependent polarity propagation can operate across discontinuous neuronal structures remains unclear. During pupal development of the *Drosophila* medulla, R8 photoreceptor axon terminals transiently form horseshoe-shaped structures whose orientation reflects neuronal column polarity. Here, we show that Frizzled (Fz) progressively becomes asymmetrically localized within developing horseshoe structures, whereas Van Gogh (Vang) transiently accumulates in adjacent glial cells during early developmental stages. Conditional disruption of Vang in glial cells impaired asymmetric Fz localization and disrupted horseshoe orientation polarity. Glial Vang localization preceded the emergence of robust Fz asymmetry, suggesting that glial Vang functions during an early polarity establishment phase. Furthermore, mosaic knockdown of *fz* in R8 neurons altered Fz localization in neighboring R8 terminals, suggesting local propagation of polarity information between adjacent neuronal columns. Together, our results suggest that PCP-dependent polarity propagation operates across discontinuous neuronal columns in the developing medulla and provide a framework for understanding coordinated polarity formation in neural tissues lacking continuous epithelial organization.

## Introduction

Neural circuits are organized into highly ordered structural units that enable efficient information processing. Across animals ranging from mammals to insects, neurons are arranged into columnar structures in which distinct neuronal populations are spatially aligned and functionally coordinated [1–3]. In the mammalian cerebral cortex, neurons form radially organized cortical columns that serve as fundamental units of sensory and cognitive processing [3]. Despite the importance of these structures, the molecular mechanisms that establish polarity and directional organization within neuronal columns remain poorly understood.

The *Drosophila* visual system provides a powerful model for investigating neuronal column organization [2]. In the medulla neuropil, approximately 800 columns are arranged in a regular hexagonal lattice along the dorsal–ventral axis [1]. Each column contains photoreceptor axons and multiple classes of interneurons that form stereotyped synaptic connections [2]. Previous studies demonstrated that medulla columns self-organize through regulated neuronal interactions [4]. During pupal development, R8 photoreceptor axons transiently pause within the superficial medulla layers (M1–M2) and form characteristic horseshoe-shaped terminal structures [5]. Importantly, the orientation of these horseshoe structures is spatially coordinated across the medulla: horseshoes in ventral regions face dorsally, whereas horseshoes in dorsal regions face ventrally. This stereotyped organization provides a robust morphological readout of medulla column polarity.

Planar cell polarity (PCP) signaling coordinates tissue-scale polarity in epithelial tissues through asymmetric localization of core PCP proteins, including Frizzled (Fz), Van Gogh (Vang), and Flamingo (Fmi) [6–8]. In canonical epithelial PCP systems, neighboring cells exchange polarity information through intercellular PCP complexes, allowing local asymmetries to propagate across continuous epithelial sheets and generate coherent tissue-wide polarity [9,10]. These mechanisms have been extensively characterized and reviewed in epithelial systems such as the *Drosophila* wing and eye [11,12].

Recent studies suggested that PCP factors also contribute to neuronal column organization in the *Drosophila* medulla [5]. Knockdown of *fz* or *fz2* disrupts the orientation of R8 horseshoe structures, indicating that PCP signaling contributes to polarity formation within neuronal columns [5]. In addition, PCP signaling has been implicated in multiple aspects of neural development, including axon guidance and growth cone regulation [13]. Glial cells have also been implicated in photoreceptor axon targeting through Flamingo-dependent interactions [14]. These observations raise the possibility that PCP-dependent polarity signaling also contributes to neuronal polarity organization in the medulla.

In this study, we investigated how PCP factors regulate polarity formation in developing R8 terminals. We found that Fz progressively becomes asymmetrically localized within individual horseshoe structures during pupal development, whereas Vang transiently accumulates in adjacent glial cells during early developmental stages. Conditional disruption of *vang* in glial cells impaired asymmetric Fz localization and disrupted horseshoe orientation polarity, suggesting that glial Vang functions during an early polarity establishment phase. Furthermore, mosaic knockdown of *fz* in R8 neurons altered Fz localization in neighboring R8 terminals, suggesting that polarity information propagates locally between adjacent neuronal columns. Together, our results support a model in which PCP-dependent polarity propagation contributes to coordinated polarity formation across discontinuous neuronal columns in the developing *Drosophila* medulla.

## Results

### R8 photoreceptor terminals exhibit spatially coordinated horseshoe polarity in the developing medulla

The *Drosophila* medulla is composed of regularly arranged neuronal columns that receive stereotyped photoreceptor inputs [1,2]. Each medulla column contains photoreceptor axons and multiple classes of interneurons that form highly ordered synaptic connections [1,2]. R7 photoreceptors express the ultraviolet-sensitive rhodopsins Rh3 and Rh4, whereas R8 photoreceptors express the blue-sensitive Rh5 and green-sensitive Rh6 rhodopsins, thereby contributing to wavelength-specific color vision [15]. During pupal development, R8 photoreceptor axons transiently pause within the superficial medulla layers (M1–M2) and form characteristic horseshoe-shaped terminal structures [5,16] (Fig. 1A,B).

**Fig 1.**
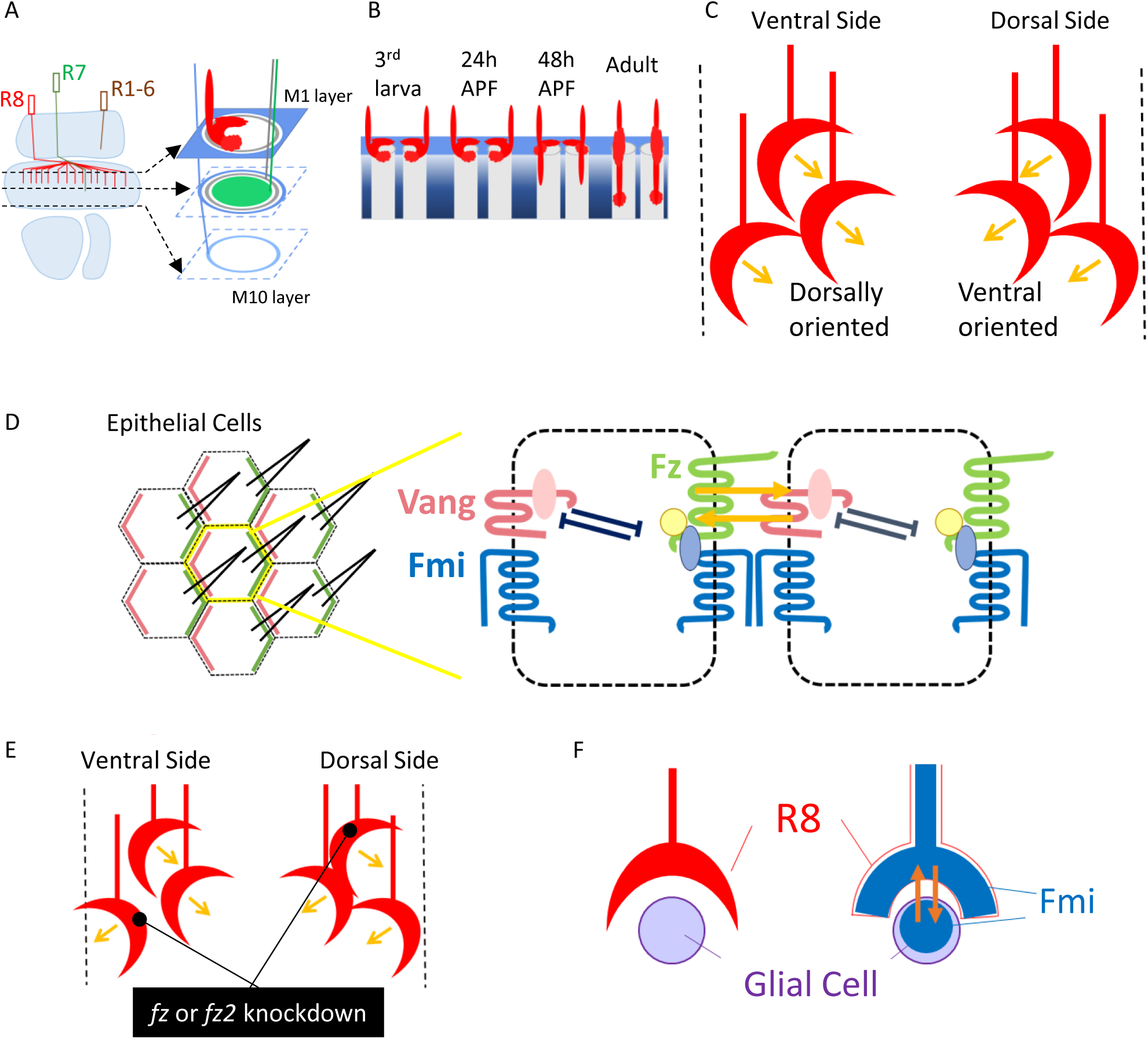
Spatially coordinated horseshoe polarity in the developing *Drosophila* medulla. (A) Schematic of the *Drosophila* optic lobe and medulla column organization. R8 photoreceptor axons project into superficial medulla layers (M1–M2), where transient horseshoe-shaped terminal structures are formed during pupal development. (B) Developmental progression of R8 terminal morphology from the third instar larval stage to adulthood. R8 terminals transiently form horseshoe-shaped structures during pupal development. (C) Schematic illustrating spatially coordinated horseshoe orientation polarity along the dorsal–ventral axis of the medulla. Horseshoes in ventral regions are dorsally oriented, whereas horseshoes in dorsal regions are ventrally oriented. (D) Simplified schematic of epithelial PCP organization and asymmetric PCP complexes [4–10,22]. Core PCP proteins, including Fz, Vang, and Fmi, form asymmetric intercellular complexes between neighboring cells. (E) Simplified schematic summarizing previous findings that knockdown of *fz* or *fz2* disrupts horseshoe orientation polarity in the developing medulla [13]. (F) Simplified schematic model based on previous studies proposing Flamingo-dependent interactions between glial cells and developing R8 terminals during horseshoe formation [14].

Importantly, these horseshoe structures exhibit coordinated orientation polarity along the dorsal–ventral axis of the medulla[5]. Horseshoes located in ventral regions are oriented dorsally, whereas horseshoes located in dorsal regions are oriented ventrally (Fig. 1C). This stereotyped organization provides a robust morphological readout of medulla column polarity.

PCP signaling establishes coordinated tissue polarity in epithelial tissues through asymmetric localization of core PCP proteins, including Fz, Vang, and Fmi, and through propagation of polarity information between neighboring cells [6–12]. Previous studies demonstrated that disruption of PCP factors such as *fz* and *fz2* impairs the orientation of R8 horseshoe structures in the developing medulla [5] (Fig. 1D,E), suggesting that PCP signaling contributes to neuronal column organization. In addition, glial cells interact closely with developing R8 terminals during horseshoe formation [14] (Fig. 1F), raising the possibility that glia-associated PCP signaling contributes to polarity formation within medulla columns.

However, unlike canonical epithelial PCP systems, neighboring R8 terminals in the medulla are spatially separated and do not form a continuous epithelial sheet. How polarity information is established and coordinated across such discontinuous neuronal columns therefore remains unclear.

### Frizzled becomes asymmetrically localized within developing R8 horseshoe terminals

To investigate how PCP signaling regulates polarity formation within developing medulla columns, we examined the localization dynamics of Fz during R8 horseshoe development. To visualize endogenous Fz in genetically defined cells, we generated a split-GFP knock-in reporter in which Fz was tagged with seven tandem GFP11 repeats, enabling Fz visualization in cells expressing the complementary GFP1–10 fragment (Fig. 2A) [17–19]. GFP fluorescence was reconstituted specifically in R8 photoreceptor neurons by expressing GFP1–10 under the control of the GMR driver (Fig. 2A).

**Fig 2.**
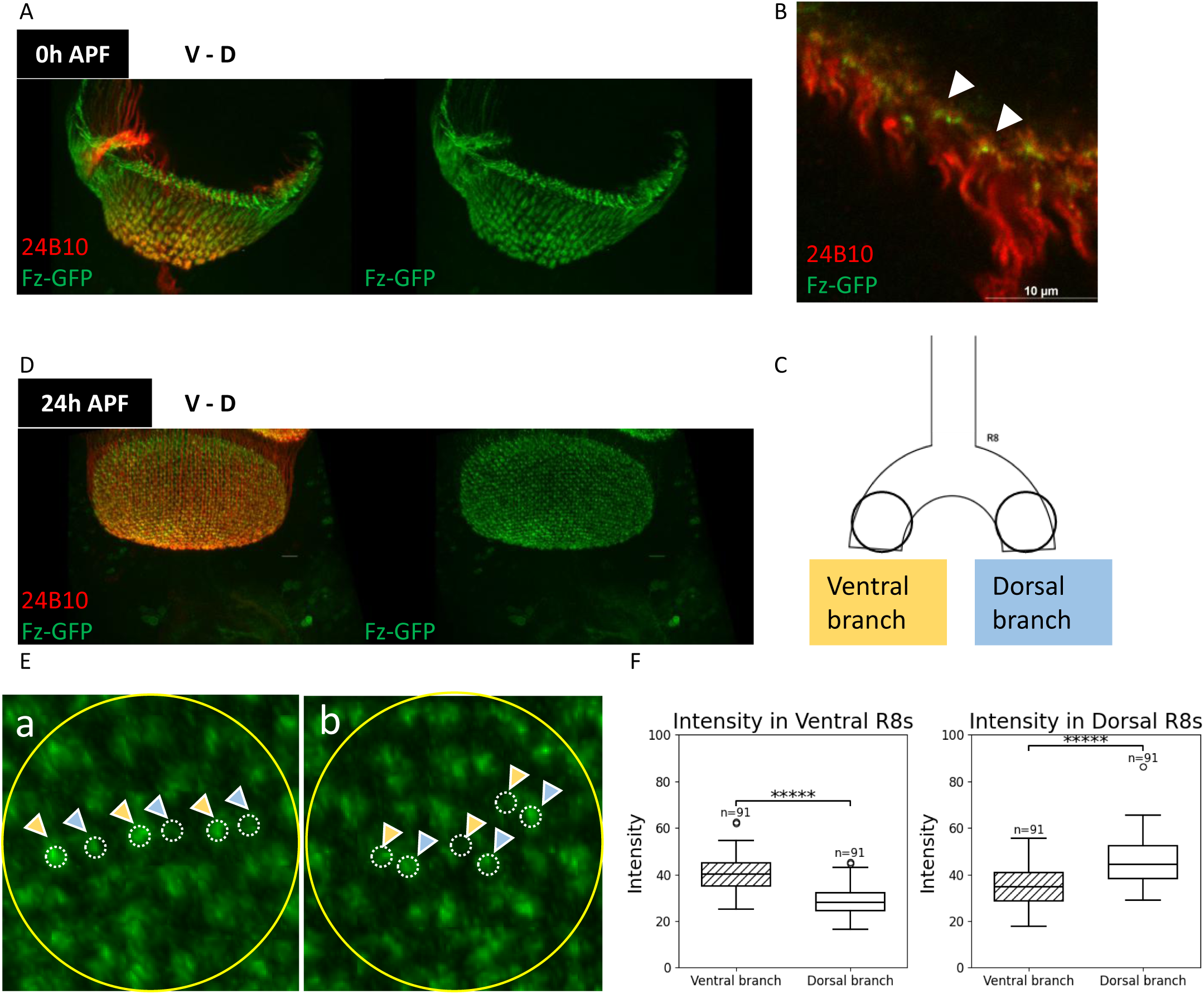
Frizzled becomes asymmetrically localized in developing R8 terminals. (A) Whole-medulla view of Fz-GFP localization at 0 h APF. Left: merged image. Right: Fz-GFP channel only. (B) Higher magnification image of Fz-GFP localization at 0 h APF. Arrowheads indicate Fz accumulation near the center of developing horseshoe structures. (C) Schematic defining dorsal and ventral branches within individual horseshoe structures for quantitative analysis. (D) Whole-medulla view of Fz-GFP localization at 24 h APF. Left: merged image. Right: Fz-GFP channel only. (E) Higher magnification images of ventrally oriented (left) and dorsally oriented (right) horseshoe structures at 24 h APF. Arrowheads indicate asymmetric Fz localization within individual branches. (F) Quantification of Fz-GFP fluorescence intensity in dorsal and ventral branches. Fz asymmetry correlates with horseshoe orientation polarity. ****p < 0.0001.

At 0 h after puparium formation (APF), Fz-GFP accumulated near the center of developing horseshoe structures without obvious branch-specific asymmetry (Fig. 2B). By contrast, at 24 h APF, Fz-GFP exhibited clear asymmetric localization within individual horseshoe structures within superficial medulla layers of the medulla (Fig. 2D). Higher magnification analysis revealed that Fz preferentially accumulated on one branch of each horseshoe structure depending on its orientation along the dorsal–ventral axis (Fig. 2F).

To quantitatively evaluate this asymmetry, we defined dorsal and ventral branches within individual horseshoe structures and measured Fz-GFP fluorescence intensity in each region (Fig. 2C). In ventrally positioned horseshoes, Fz-GFP preferentially accumulated in ventral branches, whereas dorsally positioned horseshoes exhibited stronger Fz localization in dorsal branches (Fig. 2F). These results indicate that Fz progressively acquires asymmetric localization during the establishment of R8 horseshoe polarity.

### Vang transiently localizes in glial cells adjacent to developing R8 terminals

To examine Vang localization during horseshoe polarity formation, we generated a split-GFP knock-in reporter in which endogenous Vang was tagged with seven tandem GFP11 repeats. GFP fluorescence was reconstituted in glial cells by expressing GFP1–10 using the glial driver *loco*-GAL4 (Fig. 3A).

**Fig 3.**
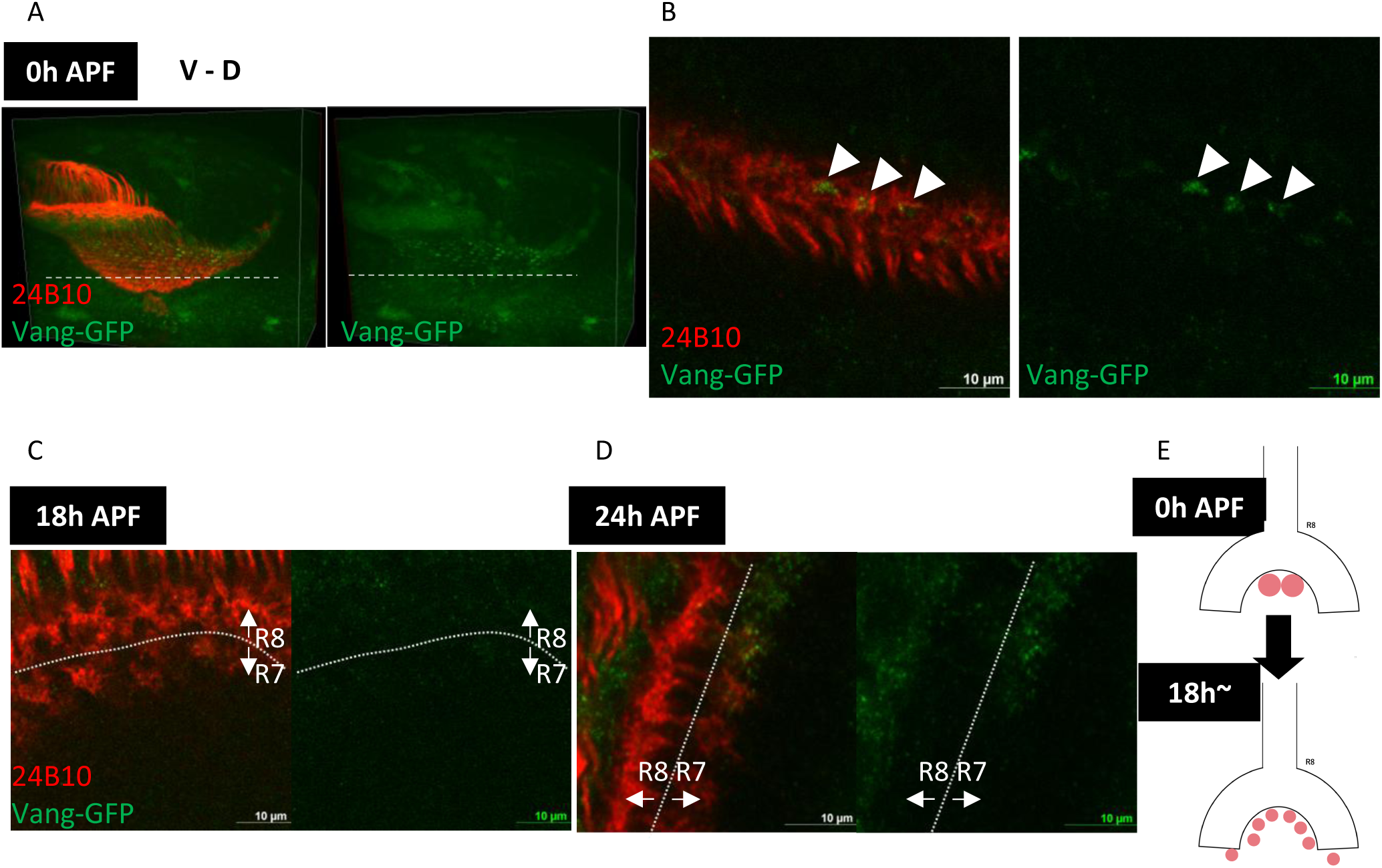
Glial Vang transiently accumulates during early stages of R8 polarity formation. (A) Whole-medulla view of glial Vang-GFP localization at 0 h APF. Left: merged image. Right: Vang-GFP channel only. (B) Higher magnification images of glial Vang localization at 0 h APF. Arrowheads indicate punctate Vang-GFP signals adjacent to developing R8 terminals. (C) Glial Vang-GFP localization at 18 h APF. Vang signals are preferentially retained in posterior regions corresponding to younger medulla columns. (D) Glial Vang-GFP localization at 24 h APF. Vang signals are substantially reduced compared with earlier developmental stages. (E) Model illustrating transient glial localization of Vang during early stages of horseshoe polarity formation.

At 0 h APF, strong Vang-GFP signals were detected throughout the superficial medulla in glial cells adjacent to developing R8 terminals (Fig. 3A,B). High-magnification analysis revealed punctate Vang signals positioned near the opening region of developing horseshoe structures (Fig. 3B).

At later developmental stages, glial Vang localization progressively decreased. At 18 h APF, Vang-GFP signals remained detectable preferentially in posterior regions corresponding to younger medulla columns (Fig. 3C). By 24 h APF, Vang-GFP localization near R8 terminals was substantially reduced (Fig. 3D). These observations indicate that glial Vang transiently accumulates adjacent to developing R8 terminals during the early stages of horseshoe polarity formation.

### Glial-specific knockout of *vang* impairs asymmetric Fz localization in R8 terminals

To investigate the role of glial Vang in R8 polarity formation, we conditionally disrupted *vang* in glial cells using a CRISPR/Cas9-based approach (Fig. 4A). Glial Vang-GFP signals observed at 0 h APF (Fig. 3A) were markedly reduced following glial-specific knockout of *vang* (Fig. 4B), indicating efficient disruption of *vang* in glial cells.

**Fig 4.**
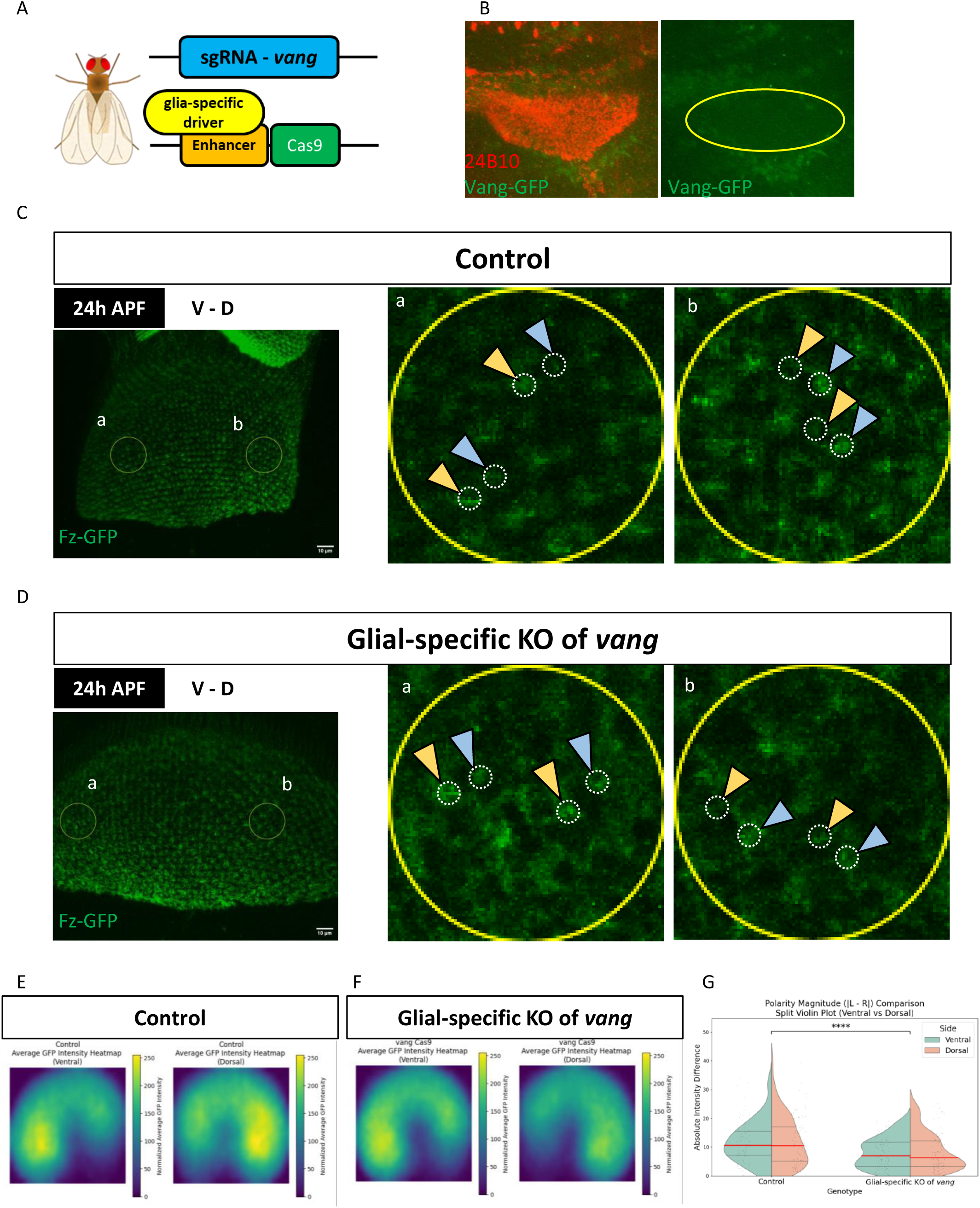
Glial-specific knockout of *vang* impairs asymmetric Fz localization in R8 terminals. (A) Schematic of the glial-specific CRISPR/Cas9-mediated *vang* disruption strategy. (B) Vang-GFP localization following glial-specific knockout of *vang*. Vang signals are strongly reduced in glial regions. (C) Fz-GFP localization in control flies at 24 h APF. Insets show representative ventrally oriented and dorsally oriented horseshoe structures exhibiting asymmetric Fz localization. (D) Fz-GFP localization following glial-specific knockout of *vang*. Insets show reduced asymmetric Fz localization within horseshoe structures. (E,F) Heatmap analyses of averaged Fz-GFP localization patterns in control flies (E) and glial *vang* disruption flies (F). (G) Quantification of Fz polarity magnitude in control and glial vang disruption conditions. Polarity magnitude was defined as the absolute fluorescence intensity difference between ventral and dorsal branches within individual horseshoe structures. Glial-specific knockout of *vang* significantly reduces asymmetric Fz localization (p = 1.86 × 10⁻⁷, ****p < 0.0001).

We next examined the effects of glial *vang* disruption on Fz localization within developing R8 terminals. In control flies, Fz-GFP exhibited characteristic asymmetric localization within horseshoe structures across the medulla (Fig. 4C). In contrast, glial-specific knockout of *vang* strongly impaired this asymmetric pattern and resulted in diffuse Fz localization within R8 terminals (Fig. 4D).

To quantitatively evaluate these changes, average fluorescence distributions were visualized using heatmap analyses (Fig. 4E,F). Control medullae displayed clear branch-biased Fz localization patterns, whereas glial Vang disruption substantially weakened Fz asymmetry. To quantitatively evaluate branch asymmetry, we calculated the absolute fluorescence intensity difference between ventral and dorsal branches within individual horseshoe structures as a polarity magnitude index. Quantification of this polarity magnitude revealed a significant reduction in asymmetric Fz localization following glial-specific knockout of *vang* (Fig. 4G). These results demonstrate that glial Vang is required for proper asymmetric localization of Fz during R8 horseshoe polarity formation.

### Glial *vang* loss disrupts horseshoe orientation polarity

We next investigated whether disruption of asymmetric Fz localization affects tissue-scale horseshoe polarity organization. In control flies, horseshoe structures exhibited coherent orientation polarity along the dorsal–ventral axis of the medulla (Fig. 5A). By contrast, glial-specific knockout of *vang* caused substantial disorganization of horseshoe orientation patterns (Fig. 5B).

**Fig 5.**
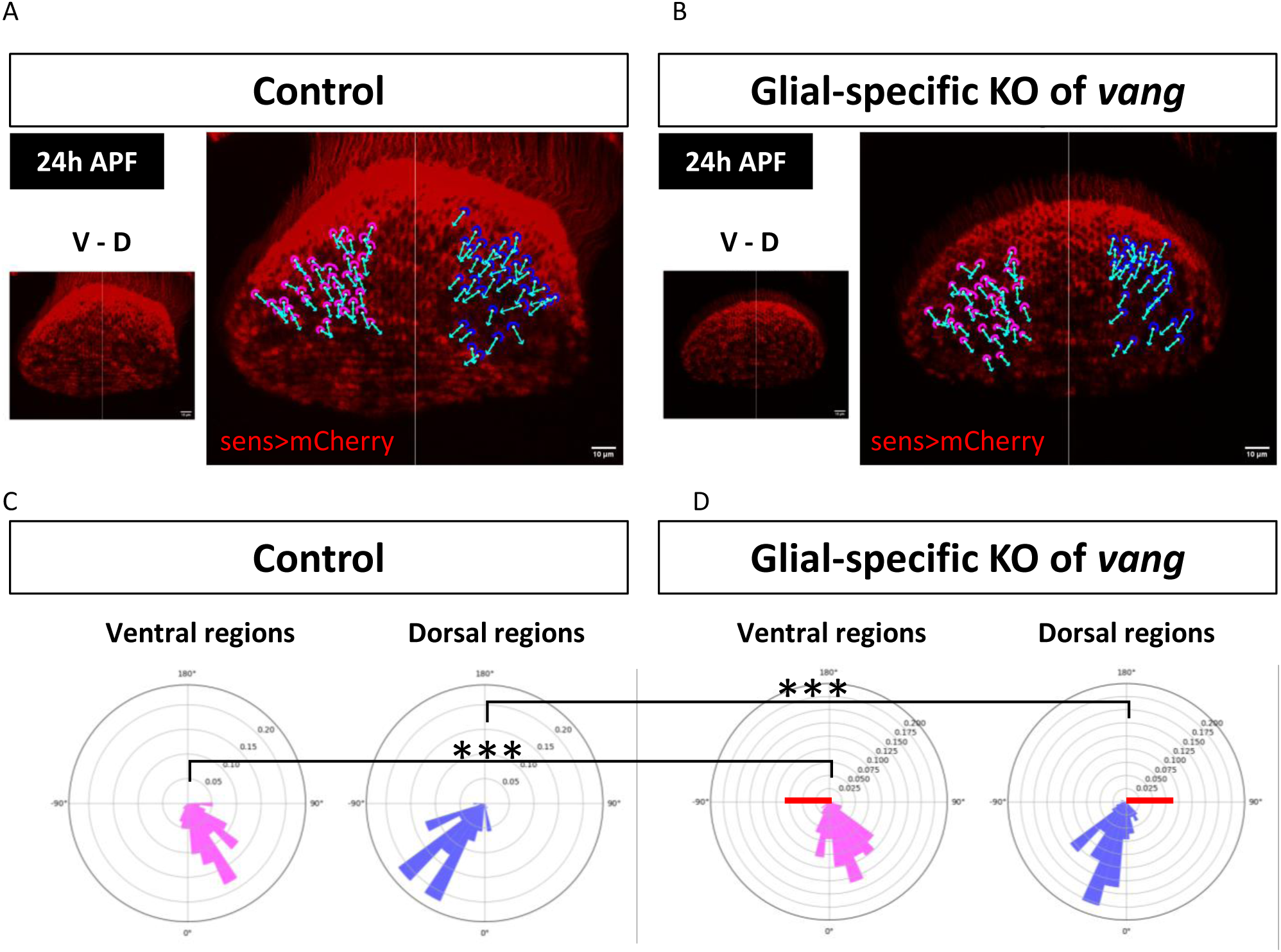
Glial *vang* knockout impairs horseshoe orientation polarity. (A) Representative annotation image showing horseshoe orientation polarity in control flies at 24 h APF. (B) Representative annotation image showing disrupted horseshoe orientation polarity following glial-specific knockout of *vang*. (C) Polar histograms showing orientation angle distributions of annotated horseshoe structures in ventral and dorsal regions of control medullae. (D) Polar histograms showing orientation angle distributions of annotated horseshoe structures in ventral and dorsal regions following glial-specific knockout of *vang*. Statistical comparisons were performed separately between control and glial-specific knockout of *vang* conditions for ventral and dorsal regions. ***p < 0.001.

To quantify these defects, orientation angles of individual horseshoe structures were measured and visualized using polar histograms (Fig. 5C,D). Control medullae displayed strong directional bias in both ventral and dorsal regions, whereas glial-specific knockout of *vang* significantly disrupted coordinated horseshoe orientation polarity on each side of the medulla. Statistical comparisons between control and glial-specific knockout of *vang* conditions were performed separately for ventral and dorsal regions.

### Mosaic knockdown of *fz* alters polarity in neighboring R8 terminals

To investigate whether polarity information propagates locally between neighboring medulla columns, we induced mosaic knockdown of *fz* in R8 photoreceptor neurons using a *sens*-FLP-based stochastic mosaic strategy. In this system, *sens*-FLP was used together with a GMR-FRT-stop-FRT-GAL4 cassette to induce mosaic GAL4 expression in subsets of developing R8 neurons, thereby driving UAS-fz RNAi expression in stochastic populations of R8 cells. This approach generated mosaic regions containing neighboring control and *fz* knockdown R8 terminals within the developing medulla.

In control mosaics, Fz-GFP exhibited characteristic asymmetric localization within individual horseshoe structures throughout the medulla (Fig. 6A). In contrast, mosaic knockdown of *fz* disrupted asymmetric Fz localization not only within *fz* knockdown R8 terminals, but also in neighboring R8 terminals adjacent to mosaic boundaries (Fig. 6B).

**Fig 6.**
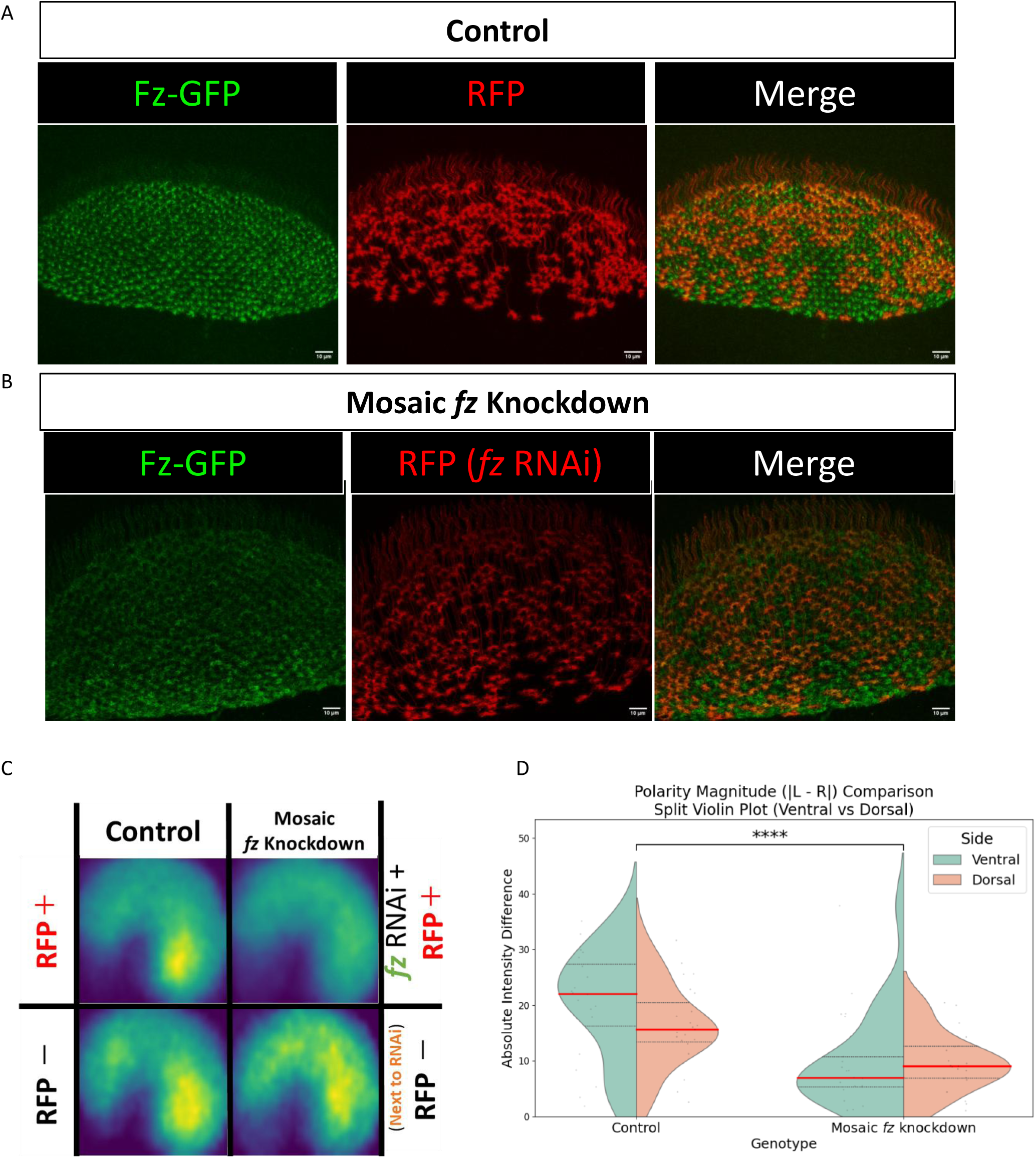
Mosaic knockdown of *fz* alters polarity in neighboring R8 terminals. (A) Fz-GFP localization in control mosaic medullae at 24 h APF. Left: Fz-GFP channel. Middle: RFP channel. Right: merged image. (B) Fz-GFP localization following mosaic knockdown of *fz* in R8 neurons. Left: Fz-GFP channel. Middle: RFP-labeled *fz* RNAi cells. Right: merged image. (C) Heatmap analyses of averaged Fz localization patterns in control mosaics and mosaic *fz* knockdown conditions. Top: RFP-positive cells. Bottom: neighboring RFP-negative cells adjacent to *fz* RNAi cells. (D) Quantification of branch-specific Fz polarity magnitude. Polarity magnitude was defined as the absolute fluorescence intensity difference between ventral and dorsal branches within individual horseshoe structures (p = 4.13 × 10⁻⁶, ****p < 0.0001).

To quantitatively evaluate these effects, average fluorescence distributions were visualized using heatmap analyses (Fig. 6C). Neighboring regions adjacent to *fz* knockdown mosaics exhibited altered Fz localization patterns compared with control regions (Fig. 6C). To quantitatively evaluate these changes, we calculated the absolute fluorescence intensity difference between ventral and dorsal branches within individual horseshoe structures as a polarity magnitude index. Quantification of this polarity magnitude demonstrated a significant reduction in Fz asymmetry in neighboring R8 terminals surrounding mosaic knockdown regions (Fig. 6D).

These observations suggest that polarity information propagates locally between adjacent neuronal columns during R8 horseshoe polarity formation in the developing medulla.

## Discussion

In this study, we investigated how PCP signaling contributes to neuronal column polarity formation in the *Drosophila* medulla. We found that Fz becomes asymmetrically localized within developing R8 horseshoe structures (Fig. 2E), whereas Vang transiently localizes in adjacent glial cells rather than in R8 neurons themselves (Fig. 3B). Furthermore, glial-specific knockout of *vang* impaired both asymmetric Fz localization and horseshoe orientation polarity (Fig. 4, Fig. 5). Finally, mosaic knockdown of *fz* in R8 neurons altered Fz localization in neighboring R8 terminals, suggesting that polarity information propagates locally between adjacent neuronal columns (Fig. 6). Together, these findings support a model in which PCP-dependent polarity propagation contributes to coordinated polarity formation across discontinuous neuronal columns.

PCP signaling has long been understood as a central mechanism underlying tissue-scale polarity formation in epithelial systems [6–12]. In canonical epithelial PCP systems, including the *Drosophila* wing and eye, core PCP proteins such as Fz, Vang, and Fmi form asymmetric intercellular complexes between neighboring cells, allowing local polarity information to propagate across continuous epithelial sheets and generate coherent tissue-wide polarity [3,4]. Although PCP signaling has also been implicated in various aspects of neural development, including axon guidance and dendritic morphogenesis, its role in neuronal column polarity formation and polarity propagation mechanisms has remained poorly understood.

Recent work by Han et al. demonstrated that horseshoe orientation polarity in the *Drosophila* medulla is disrupted by knockdown of *fz* and *fz2*, providing the first evidence that PCP signaling contributes to neuronal column organization [5]. However, the cellular basis of PCP signaling in the medulla and the mechanisms underlying polarity establishment and propagation remained unknown. By identifying asymmetric Fz localization and transient glial Vang localization, our study provides a cellular framework for PCP-dependent polarity formation in developing medulla columns.

One of the most striking findings of this study is that Vang localized transiently in glial cells rather than within R8 neurons themselves. In canonical epithelial PCP systems, Fz and Vang are asymmetrically localized in neighboring epithelial cells and exchange polarity information directly across cell-cell interfaces. In contrast, our results suggest that polarity establishment in the medulla involves glia–neuron interactions, in which glial Vang contributes to the establishment of asymmetric Fz localization within adjacent R8 terminals (Fig. 7).

**Fig 7.**
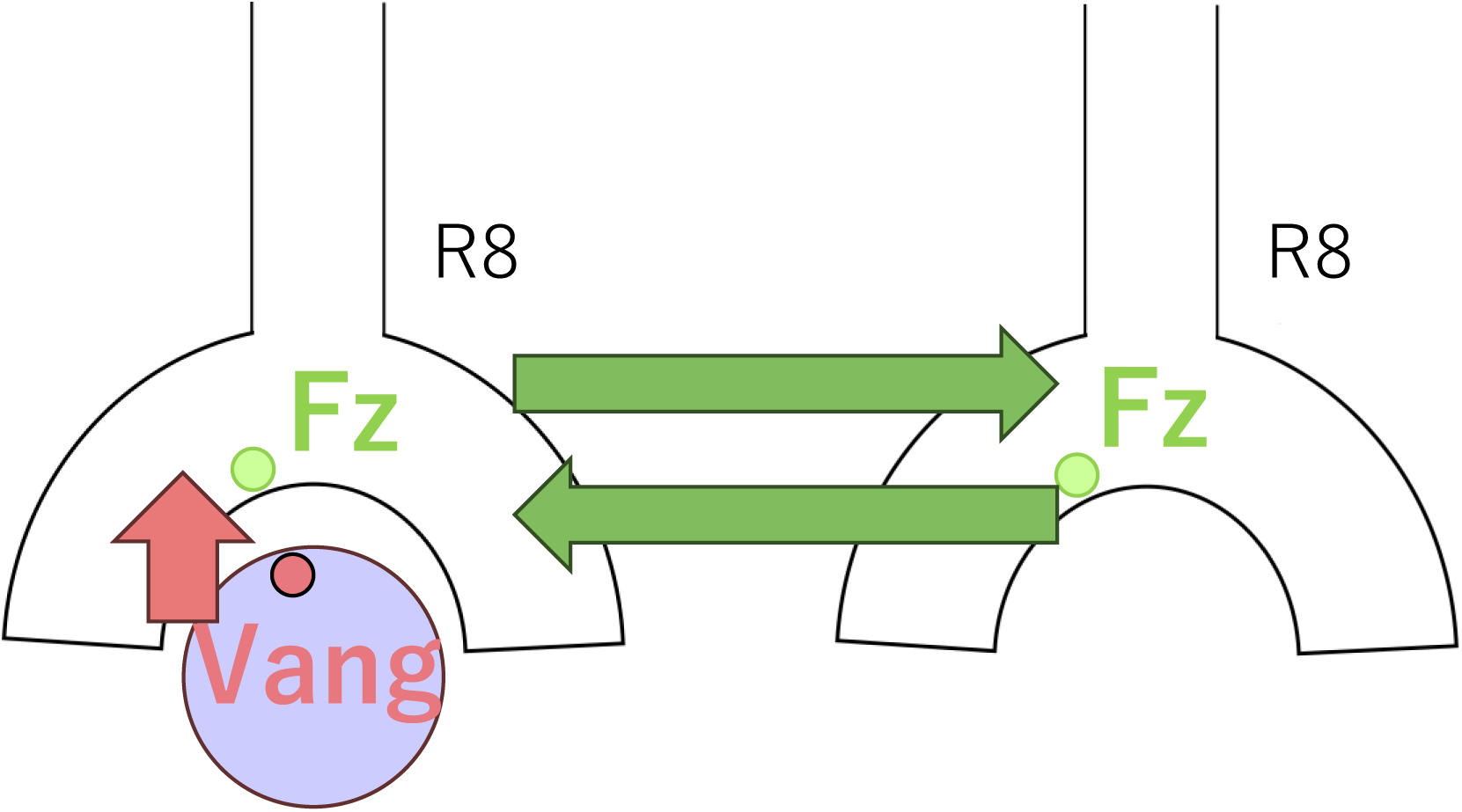
Model for PCP-dependent polarity propagation across discontinuous neuronal columns. Proposed model illustrating PCP-dependent polarity establishment and local polarity propagation in the developing medulla. Transient glial Vang localization contributes to the establishment of asymmetric Fz localization within developing R8 terminals during early developmental stages. Local polarity information is subsequently propagated between neighboring R8 columns, resulting in coordinated horseshoe orientation polarity across the medulla.

Consistent with this model, glial-specific knockout of *vang* impaired asymmetric Fz localization and disrupted horseshoe orientation polarity. Importantly, glial Vang localization was strongest at 0 h APF and progressively decreased during subsequent developmental stages, whereas robust Fz asymmetry became clearly detectable by 24 h APF. These temporal dynamics suggest that glial Vang functions during an early polarity establishment phase, after which polarity states may become stabilized or reinforced within developing neuronal columns.

Our mosaic *fz* knockdown experiments further suggest that polarity information propagates locally between neighboring R8 columns. In canonical epithelial PCP systems, polarity propagation occurs through direct contact between adjacent epithelial cells. However, neighboring R8 terminals in the medulla are spatially separated and do not form a continuous epithelial sheet. Thus, our findings suggest that PCP-dependent polarity propagation can occur even across discontinuous neuronal columns lacking canonical epithelial organization.

How polarity information propagates between discontinuous neuronal columns remains unclear. One possible mechanism is that transient filopodia-like protrusions extending from developing R8 growth cones mediate local inter-column interactions. Developing photoreceptor axons dynamically extend protrusions during medulla targeting [20], raising the possibility that transient contacts between neighboring columns facilitate local polarity alignment. However, the present study does not directly address this mechanism, and future analyses using live imaging or super-resolution microscopy will be required to test this hypothesis.

Another important unresolved question concerns the origin of the global polarity axis within the medulla. Previous studies suggested that the Wnt family ligand DWnt4 contributes to medulla polarity formation [2]. It is therefore possible that a DWnt4-dependent global directional cue biases initial PCP orientation, which is subsequently amplified through local PCP-dependent polarity propagation between neighboring columns.

Together, our findings suggest that PCP-dependent polarity propagation operates across discontinuous neuronal columns in the developing *Drosophila* medulla. More broadly, our findings suggest that PCP-dependent polarity propagation may represent a general strategy for coordinating polarity organization in neural tissues lacking continuous epithelial architecture.

## Materials and methods

### Fly husbandry

Flies were maintained on standard cornmeal medium at 25°C unless otherwise noted. All experiments in this study were performed using animals raised at 25°C.

### Fly stocks and genetics

The following GAL4 drivers were used in this study: GMR-GAL4 for photoreceptor-specific expression and *loco*-GAL4 (BDSC #602886) for glial-specific expression. Mosaic knockdown experiments were performed using a *sens*-FLP (BDSC #55796)-based stochastic recombination system in combination with an FRT-stop-FRT-GAL4 (kindly provided by M. Sato, Kanazawa University) cassette.

For visualization of endogenous PCP proteins, CRISPR/Cas9-mediated GFP11x7 knock-in lines for *fz* and *vang* were used (this work) [20]. GFP fluorescence was reconstituted by expressing UAS-splitGFP1-10 (kindly provided by M. Sato, Kanazawa University) or GMR-splitGFP1-10 (internal stock) in specific cell types using GAL4 drivers. For R8-specific visualization, *sens*-GAL4 [21] was used to drive GFP1–10 expression.

Glial-specific knockout of *vang* was achieved using a CRISPR/Cas9-based conditional knockout strategy. A ubiquitously expressed *vang* sgRNA line (BDSC #77214) was combined with UAS-Cas9 (BDSC #54592) and *loco*-GAL4 to induce glial-specific knockout of *vang*. The *sens-lexA* and lexAop-mCherry lines used for cell-specific labeling were kindly provided by the Sato lab (Kanazawa University).

For RNAi experiments, UAS-*fz* RNAi (BDSC #34321) and UAS-*vang* RNAi (BDSC #34354) lines were used.

### Pupal staging

White pupae were collected as 0 h APF. For analysis of developmental stages, animals were incubated at 25°C for the indicated periods after puparium formation. For 24 h APF samples, brains were dissected 22–24 h after white pupal collection.

### Immunohistochemistry

Pupal brains were dissected in 0.1% PBT (PBS containing 0.1% Triton X-100) under a stereomicroscope. Dissected brains were fixed in 4% paraformaldehyde in 0.1% PBT for 50 min at 25°C with gentle agitation.

After fixation, samples were washed three times with 0.1% PBT and incubated with primary antibodies diluted in 0.1% PBT overnight or for two nights at 4°C. Samples were subsequently washed three times with 0.1% PBT and incubated with secondary antibodies diluted in 0.1% PBT for 1 h at 25°C. After secondary antibody incubation, brains were washed three times with 0.1% PBT, rinsed once with PBS, and mounted in VECTASHIELD mounting medium (Vector Laboratories).

### Confocal imaging

Images were acquired using a Nikon A1 confocal laser scanning microscope equipped with a Nikon Ni-E microscope system. A 40× objective lens was used for all imaging experiments.

For whole-medulla imaging, z-stack images were acquired at 1 μm intervals spanning the entire region containing R8 terminals. For imaging of individual horseshoe structures, z-stack images were acquired at 0.5 μm intervals over a depth of 2–6 μm.

Maximum intensity projection images were generated using NIS-Elements AR Analysis software (Nikon).

### Quantification of Fz asymmetry

For quantitative analysis of Fz localization, dorsal and ventral branches within individual horseshoe structures were manually defined as regions of interest (ROIs). Mean fluorescence intensity within each ROI was measured using Adobe Photoshop (Adobe).

For heatmap analyses, horseshoe structures were manually annotated and spatial coordinates were integrated across samples using a custom Python-based analysis pipeline. Spatial probability distributions were visualized as averaged heatmaps. Analysis scripts are publicly available on GitHub (https://github.com/morita761/horseshoe-heatmap-analyzer).

### Horseshoe orientation analysis

Orientation polarity of horseshoe structures was analyzed using a custom Python-based analysis pipeline. Orientation angles were calculated based on the opening direction of individual horseshoe structures, and angular distributions were visualized as polar histograms. Analysis scripts are available on GitHub (https://github.com/morita761/arrow-orientation-analyzer).

### Statistical analysis

Statistical analyses were performed using SciPy and Statsmodels libraries in Python (Python version 3.11.9).

For comparisons between two groups, Welch’s t-test was used. For experiments involving multiple variables, two-way ANOVA followed by Tukey’s honestly significant difference (HSD) post hoc test was performed.

Statistical significance was indicated as follows: n.s., not significant (p ≥ 0.05); *p < 0.05; **p < 0.01; ***p < 0.001; ****p < 0.0001.

## Supporting information

List of Genotypes

## Acknowledgments

We are grateful to the Bloomington Drosophila Stock Center (BDSC) for providing fly stocks used in this work. We would like to express our gratitude to WellGenetics Inc. for their professional assistance in generating fly constructs, and to Eurofins Genomics for plasmid synthesis and preparation.

## Author Contributions

T.M. performed experiments, developed analysis software, analyzed data, prepared figures, and wrote the manuscript.

H.Y. performed characterization of Fz and Vang localization during medulla development.

M.S. provided materials and techniques, and helped statistic analysis.

J.O. supervised the study and edited the manuscript.

T.S. conceived, supervised the study and edited the manuscript.

## Supporting tables

**S1 Table. The list of genotypes of flies used in this study.**

